# A Multiparametric Analysis Reveals Differential Behavior of Spheroid Cultures on Distinct Ultra-Low Attachment Plates Types

**DOI:** 10.1101/2024.03.26.586778

**Authors:** Mario Vitacolonna, Roman Bruch, Ane Agaçi, Elina Nürnberg, Tiziana Cesetti, Florian Keller, Francesco Padovani, Simeon Sauer, Kurt M. Schmoller, Markus Reischl, Mathias Hafner, Rüdiger Rudolf

## Abstract

Spheroids have become principal three-dimensional biological models to study cancer, developmental processes, and drug efficacy. For spheroid generation, ultra-low attachment plates are noteworthy due to their simplicity, compatibility with automation, and experimental and commercial accessibility. Nonetheless, it is unknown whether and to what degree the plate type impacts spheroid formation and biology. This study employed automated brightfield microscopy to systematically compare the size and eccentricity of spheroids formed in six different plate types using four distinct human cell lines, i.e., CCD-1137Sk fibroblasts, HaCaT keratinocytes, and MDA-MB-231 and HT-29 cancer cells. Results showed that all plate types exhibited similar sphe-roid-forming capabilities, and the gross patterns of growth or shrinkage during four days after seeding were comparable. Yet, size and eccentricity varied systematically among specific cell lines and plate types. A confocal wholemount analysis by a novel pipeline of AI-based 3D-image analysis procedures revealed changes in cell proliferation, cell number, nuclear volume, and keratinocyte differentiation, which were accompanied by altered YAP1-signals. The findings show that the plate type may influence the outcome of experimental campaigns. It is advisable to scan different plate types for the optimal configuration for a specific investigation instead of using one standard plate for all kinds of applications.

## 1. Introduction

Three-dimensional cell cultures, such as spheroids and organoids, provide intermediate complexity and relevance as biological model systems for fundamental and applied research inquiries **[1]**. Compared to classical adherent monolayer cell cultures, 3D models are more laborious concerning handling, data acquisition, and data analysis. However, the three-dimensional approach can better reflect naturally occurring gradients of drugs, waste, nutrients, and gases than two-dimensional cultures and it usually better allows assessing the effects of extracellular matrix, cellular interactions, and drugs **[2,3]**. Concerning animal models, 3D-cell cultures are still inferior in terms of complexity and the faithful recapitulation of systemic properties, such as drug-induced release of hormones or activation of neural or immune responses. Yet, lab-on-a-chip approaches are increasingly coping even with systemic aspects **[4,5]**, organoids and assembloids are reflecting a growing number of tissue-specific characteristics **[6–8]**, and a clear advantage of the cellular models is that these can employ cells of a matched species origin, e.g., human cells for human disease studies, and even using patient-derived cells for personalized questions **[7,9]**.

Among the currently available 3D-culture models, spheroids have likely been most frequently used due to their relative ease of production and their reproducibility in terms of key features, such as size and response to drugs **[10]**. Spheroids are mostly made of immortal cell lines and are composed of a single or a few different cell types, depending on the addressed question. Spheroid production may employ scaffolding substrates, such as collagen or Matrigel, or may be achieved in a scaffold-free manner **[10–14]**. Typically, regardless of the use of scaffolds or not, the creation of spheroids is based on avoiding the attachment of cells to any other surface than neighboring cells. Therefore, techniques such as hanging drop, bioreactor culture, matrix encapsulation, magnetic levitation, or ultra-low attachment (ULA) plates have been devised **[10]**.

The high technical fidelity, that can normally be obtained with spheroid cell cultures allows for determining even subtle effects and/or testing several experimental conditions with acceptable efforts in workload, time, and materials. To achieve maximal technical robustness, all components in the testing pipeline need to perform optimally and reliably. Previous studies addressed the effects of different spheroid-generation types **[15,16]**, substrates **[17]**, surfaces **[18]**, use of microfluidics **[18]**, and media volume **[19]** on the robustness of spheroid formation. Due to their ease of use for multiple drug testing purposes, ULA plates have been of increasing relevance among the spheroid-generation modes. Several commercial ULA-plate products with similar base technology, using mostly 96-well and 384-well standard plate formats, are currently available. Systematically, this study compared 96-well ULA plates from six principal commercial sources concerning their performance in terms of spheroid formation, roundness, compactness, and growth of four human cell lines with different basic characteristics. In detail, the non-neoplastic foreskin fibroblast cell line, CCD-1137Sk, and the keratinocyte cell line, HaCaT, were used as models with low proliferative activity in 3D **[20]**. HaCaT cells were also selected, because of their capacity to display cellular differentiation **[20]**. Further, colon cancer cells, HT-29, and breast cancer cells, MDA-MB-231, were used as representatives for highly proliferative cell types, that are either easily forming spheroids in the absence of any scaffold (HT-29)**[21,22]** or that are dependent on the addition of ECM-components for efficient spheroid assembly (MDA-MB-231)[12,14,23].

Due to the known difference between adherent HaCaT and HT-29 cells in handling of the transcription factor, YAP **[24]**, spheroids of these cell lines were further analyzed with respect to total cell number and proliferation as well as expression and nuclear to cytoplasm ratio of YAP. Together with its paralog, TAZ, YAP is a principal downstream effector of the Hippo pathway, which controls cell proliferation and survival, metabolism and motility, as well as cell fate and differentiation as a function of mechanical signals **[25]**. Factors including cell-cell contacts, ECM stiffness, cell shape and stretching **[25]** control Hippo activity, whereby the downstream kinases, LATS1 and LATS2, mediate YAP phosphorylation, cytoplasmic sequestration, and inactivation of YAP-dependent gene expression. Hippo signaling is critically altered in several imbalance and disease states, such as in wound healing and cancer **[26]** and it has been intensely addressed as a drug target **[26]**. Yet, due to its pleiotropic regulation, YAP activity and function were found to be significantly different between adherent and 3D-cell cultures **[27–29]**, arguing for further research in that direction. Since YAP nuclear translocation is a major proxy for YAP activity **[25,30]**, determination of the YAP nuclear-to-cytoplasm ratio (N/C ratio) is key to single-cell assays in this field. However, automated analysis of this characteristic in 3D-samples has been hampered by technical issues and it has been achieved only for very small spheroids **[28]** or by manually checking a few cells in larger spheroids **[29]**. Thus, to automate detection of cell nuclei and YAP distribution in spheroids of several thousand cells, a novel pipeline of handling spheroids for detailed 3D-morphometric single-cell analysis was needed and established in the present work. In summary, the results demonstrated that all tested plate types consistently led to the formation of spheroids, showing the maturity and reliability of this technological platform. However, it also revealed that specific features, such as the number of nuclei or proliferating cells, the cell differentiation capacity, and the activity of specific signaling pathways can vary significantly between different plate types. This underlines the need to carefully select a specific plate type and to avoid any switch in this parameter during the acquisition of complex data sets.

## 2. Materials and Methods

### 2.1. Cell culture and spheroid generation

To investigate the influence of the cell culture plates on the generation and growth of spheroids, 96-well ULA plates from 6 different manufacturers were tested (in the following referred to as A-F): A: BIOFLOAT™ 96-well plates (faCellitate, #F202003), B: BRANDplates® 96-well microtitration plate (BrandTech Scientific, #781900), C: Cellstar® 96-well Microplate (Greiner, #650970), D: CellCarrier Spheroid ULA 96-well Microplates (PerkinElmer, #6055330), E: Corning® Costar ® 96-well Clear Round Bottom Ultra-Low Attachment Microplates (Corning, #7007), F: 96-well plate Sphera™ Low-Attachment Surface (Thermo Scientific, #174927). The following four cell lines were used to generate spheroids: CCD-1137Sk human foreskin fibroblast cells (ATCC) were cultured in Iscove’s modified Dulbecco’s medium (IMDM, Capricorn) supplemented with 10 % fetal bovine serum (FBS, Capricorn) and 1 % penicillin/streptomycin (Pen/Strep, Sigma-Aldrich). For spheroid monoculture generation, cells were detached using Trypsin/EDTA (Sigma-Aldrich) and seeded onto 96-well ULA plates at a concentration of 2 × 10^3^ cells per well. The human keratinocyte cell line HaCaT (kindly provided by BRAIN AG, Zwingenberg) was cultured in Dulbecco’s Modified Eagle Medium (DMEM) High Glucose with L-Glutamine and Sodium Pyruvate (Capricorn) supplemented with 1 % Pen/Strep and 10 % FBS. For spheroid generation, cells were detached using Trypsin/EDTA and seeded onto the ULA plates at a concentration of 5 × 10^3^ cells per well. HT-29 colon cancer cells (ATCC) were cultured in McCoy’s 5A medium (Capricorn) supplemented with 10 % FBS and 1 % Pen/Strep. For spheroid generation, cells were detached using Trypsin/EDTA and seeded onto the ULA plates at a concentration of 5 × 10^2^ cells per well. MDA-MB-231 cells were cultured in RPMI 1640 medium with L-glutamine (Capricorn) supplemented with 10% FBS and 1 % Pen/Strep. For spheroid generation, cells were detached using Trypsin/EDTA, supplemented with 5 mM type 1 collagen (Roche Diagnostics) to allow spheroid formation and seeded onto the ULA plates at a concentration of 5 × 10^3^ cells per well. Each cell line was seeded in the appropriate plate type as replicates (n=24 per cell line), and all experiments were conducted as 3 biological replicates. All cells were maintained in a humidified incubator at 37 °C with 5 % CO^2^ fumigation.

### 2.2. Brightfield imaging and measurement of spheroid diameter, eccentricity, and area covered by dissociated cells

Plates were imaged every 24 h for 4 days using the Cytation™ 5 cell imaging multi-mode plate reader with BioSpa 8 automated incubator, using a 10x phase contrast objective and Gen5 software (all BioTek Instruments). Spheroid diameter and eccentricity were automatically quantified using MATLAB with the SpheroidSizer software **[31]**. To measure the area covered by dissociated cells around the HaCaT spheroids, we used a two-step approach due to the inhomogeneous intensity distribution of the objects to be measured. First, a pixel-based classification was performed using the machine-learning based bio-image analysis tool Ilastik (V1.4.0) **[32]**. The pixel classifier aims to learn to distinguish whether each pixel belongs to a specific object type or background, using not only the intensity information of that pixel but also the intensity information of local pixel neighbors **[33]**. Two classes (background, cells) were defined and manually labeled separately using the paintbrush tool, which was used to train Ilastik’s machine-learning algorithm to identify the objects of interest. The training was iterative, adding new pixel classifications until the probability maps were stable and adequately distinguished the object types. The training was repeated for each plate type, as the background of each plate type was different. The pixel classification workflow described here performed a semantic segmentation that divides the image into two semantic classes (foreground and background), but not into individual objects. For each pixel in the image, Ilastik estimates the probability that the pixel belongs to each of the semantic classes. The resulting probability maps were then exported as .tif files and loaded into the open-source software CellProfiler (V4.2.6) **[34]** in combination with the corresponding raw brightfield images to perform segmentation of the dissociated cells. The probability maps were converted to a single channel using the ‘ColortoGray’ module to obtain only the cell channel. The images were then smoothed with a 5-pixel wide Gaussian filter and segmented using the IdentifyPrimaryObjects module using Otsu’s method with two-class thresholding and an object diameter between 5 and 200 pixels to exclude the core spheroid. The segmented objects were then quantified using the ‘MeasureObjectSizeShape’ and ‘MeasureImageAreaOccupied’ modules. The ‘CalculateMath’ and ‘DisplayDataOnImage’ modules were used to calculate the percentage of occupied area and to create overlays.

### 2.3. Whole mount immunostaining and optical clearing

Spheroids (n=10 per group) were transferred to Eppendorf tubes, washed once with phosphate-buffered saline (PBS, Sigma Aldrich), and fixed with 4 % paraformaldehyde (PFA, Carl Roth) for 1 h at 37 °C, followed by two washes with PBS containing 1 % FBS for 5 min each. To remove traces of fixative, spheroids were quenched with 0.5 M glycine (Carl Roth) in PBS for 1 h at 37 °C with gentle shaking. Spheroids were then incubated for 30 min in a penetration buffer containing 0.2 % Triton X-100, 0.3 M glycine, and 20 % DMSO (all Carl Roth) in PBS to enhance the penetration of antibodies and nuclear stains. Spheroids were then incubated in a blocking buffer (0.2 % Triton X-100, 1 % BSA, 10 % DMSO in PBS) for 2 h at 37 °C with gentle shaking. After blocking, samples were incubated with primary antibodies overnight (ON) at 37 °C with gentle shaking. Primary antibodies were diluted in antibody buffer (0.2 % Tween 20, 10 µg/ml heparin, both Sigma-Aldrich, 1 % BSA, 5 % DMSO in PBS) at the following concentrations: anti-Ki-67 (Abcam, 1:300), anti-YAP1 (Invitrogen, 1:150). Samples were then washed 5 × for 10 minutes each in wash buffer (0.2 % Tween-20, 10 µg/mL heparin, 1 % BSA) and stained with secondary antibodies goat anti-mouse IgG (H+L) Alexa Fluor®488 (Invitrogen, 1:500), donkey anti-rabbit IgG (H+L) Alexa Fluor®555 (Invitrogen, 1:800), SiR-actin (Spirochrome, 1:1000) and DAPI (SigmaAldrich, 1:1000) ON at 37 °C in antibody buffer with gentle shaking. Samples were then washed 5 × for 10 minutes in washing buffer with gentle shaking and cleared with FUnGI clearing solution (50 % glycerol (vol/vol), 2.5 M fructose, 2.5 M urea, 10.6 mM Tris Base, 1 mM EDTA) ON as previously described [35]. Cleared samples were transferred to 18 well µ-slides (Ibidi) in the same solution and kept in the microscope room for several hours to allow for temperature adjustment.

### 2.4. Spheroid cryosectioning and staining

HaCaT spheroids were collected in an Eppendorf tube for cryosectioning. After being washed twice with PBS, they were fixed with 4 % paraformaldehyde in PBS for 30 minutes at room temperature. Following this, the spheroids were incubated in 15 % sucrose (Carl Roth) in PBS overnight at 4 °C, then in 25 % sucrose in PBS again overnight at 4 °C. Subsequently, they were embedded in Tissue-Tek Cryomolds using OCT (Leica Biosystems). 15-µm thick sections were prepared using a CM-1950 cryostat (Leica Biosystems). Cryosections were permeabilized with 0.1 % Triton X-100 (Carl Roth) in PBS, then blocked with 2 % BSA in PBS before being stained with anti-Cytokeratin 10 (Abcam; 1:1000), anti-Cytokeratin 14 (ThermoFisher, 1:1000), and anti-Involucrin (Abcam; 1:1000) ON at 4 °C. Samples were washed 3 x with PBS containing 1 % FBS, followed by secondary antibody and nuclei staining using donkey anti-rabbit IgG (H+L) Alexa Fluor®488 (1:800), anti-mouse IgG (H+L) Alexa Fluor®555 (1:800), and DAPI (1:1000) for 2 h at RT. Finally, sections were washed 3 x with PBS/1 % FBS, mounted with Mowiol (Carl Roth) and imaged using a confocal microscope (SP8, Leica).

### 2.5. Image acquisition using confocal microscopy

All 3D cultures and cryosections were imaged using an inverted Leica TCS SP8 confocal microscope (Leica Microsystems CMS, Mannheim, Germany) equipped with an HC PL APO 20×/0.75 IMM CORR objective, 488 nm, 561 nm and 633 nm lasers and Leica Application Suite X software. 3D-wholemount image stacks were acquired with comparable settings, using Immersion Type F (Leica Microsystems, RI 1.52) as immersion fluid, with a resolution of 1024 × 1024 pixels (473 × 473 nm per pixel), a z-step size of 2 µm, a laser intensity of 1-1.5 % and a gain setting of 600 to avoid overexposure of pixels. All image stacks were acquired with z-compensation to compensate for depth-dependent signal loss. Cryosections were imaged with a resolution of 1024 × 1024 pixels (473 × 473 nm per pixel), a z-step size of 1 µm, a laser intensity of 0.4-1.2 % and a gain setting of 600 to avoid overexposure of pixels.

### 2.6. 3D segmentation and image analysis

Raw confocal data were converted to multi-channel .tif files using Fiji **[36]**. 3D segmentation of nuclei and fluorescence signals of Ki-67 and plasma membrane (SiR-actin) staining was performed using Cellpose (V2.2), a deep learning-based instance segmentation tool **[37]**. For each cell type and fluorescence marker, the convolutional neural network was trained on hand-annotated ground truth datasets prepared from nuclei, Ki-67, and plasma membrane datasets to improve segmentation accuracy. To prepare the annotated training data, spheroids of each cell type were first pre-segmented using the pre-trained nuclei and cyto2 models in Cellpose, including the 2 fluorescent markers for DAPI and Ki67, as an initial step. From these, 3 patches with sizes of 32 × 128 × 128 pixels (z, y, x) were extracted for each and manually corrected using the Segmentor software **[38]**. Supervised training from scratch was performed as described in **[39]** using the command line interface for Cellpose. We varied the number of training epochs from 50 to 1,000 and tested the segmentation performance with the segmentation (SEG) and detection (DET) measures used in the cell tracking challenge **[40]**. Custom-trained models with the highest DET score were selected for the subsequent segmentation of the 3D datasets. To perform the segmentation and subsequent quantitative analysis in batch mode, we used Cell-ACDC (V1.4), an open-source graphical user interface (GUI)-based framework for cell segmentation, visualization, and data analysis that embeds various neural networks such as Cellpose **[41]**. For segmentation, the appropriate custom-trained model was loaded into Cellpose with the following parameters: ‘flow_threshold=0.4’, ‘Cellprob_threshold=-2.0’, and ‘stich_threshold=0.7’ for nuclei and Ki-67 channels, and ‘stich_threshold=0.3’ for membrane channel. The cell diameters were automatically calculated by the algorithm. The output label masks were then used for downstream analysis such as quantification of object count and volumes using the regionprops function from the scikit-image Python package built into Cell-ACDC. Segmented nuclei with a volume of less than 300 µm^3^ and greater than 3,000 µm^3^ were considered debris or segmentation errors and excluded from further analysis. Quantitative analysis of spheroid volume and density was performed using dedicated Python scripts. The nuclei segmentation results were used as a starting point. This was followed by 40 iterations of binary dilation followed by 40 iterations of binary erosion to close the holes between the nuclei without increasing the overall size of the spheroid segmentation. A structuring element with a connectivity of 1 was used. The remaining holes within the spheroid segmentation were filled. If several unconnected structures remained, only the largest was used. Spheroid density was calculated by dividing the number of nuclei inside the spheroid by the volume of the segmented spheroid. The void region inside the spheroid was defined as the volume of the segmented spheroid excluding the nuclei segmentation, i.e., the region outside the nuclei and within the spheroid segmentation. To calculate the relative number of proliferative cells, the amount of Ki67^+^ cells was divided by the total number of nuclei counted.

### 2.7. Calculation of the YAP N/C ratio

To calculate the YAP1 N/C ratio at the single-cell level, we used the “Track sub-cellular objects” tool integrated into Cell-ACDC. This tool allows for the flexible selection of the minimum percentage overlap (Intersection over Union, IoU) between nuclei and membrane segmentation masks in 3D datasets to associate the objects. For our analysis, we selected a minimum IoU of 50 % (IoU ≥ 0.5). As a result of the subsequent subtraction of the segmentation masks for membranes and nuclei, a third segmentation file was generated with the cytoplasm segmentation masks (with the same IDs of the corresponding membrane and nuclei maks). Subsequently, the new generated nuclear and cytoplasmic segmentation masks with matching ID numbers were used to measure YAP1 intensities separately in the nuclei and cytoplasm in Cell-ACDC. The amount of YAP1 was measured as the mean (the sum of all pixel intensities divided by the volume), incorporating an automatic background correction (defined as all pixels outside the detected objects). Based on these mean values, the N/C ratio was calculated for all groups. In a similar manner, the segmentation masks for Ki67 were aligned with the membrane label identifiers, enabling subsequent analyses of the N/C ratio in both Ki67^+^ and Ki67^-^ cells.

### 2.8. Spatial analysis of cell populations

A custom Python-based image analysis pipeline was developed to quantitfy cellular properties from membrane, nuclear and cytoplasmic label masks, as well as raw microscopy images. The pipeline integrated several key libraries, including NumPy for array manipulations, Pandas for data handling, scikit-image for image processing, SciPy for scientific computing, and Matplotlib and Seaborn for data visualization. For each segmented cellular component, geometric features (e.g., centroid coordinates, volume) and fluorescence intensity features, from which the YAP1 N/C ratio was calculated, were extracted. Additionally, each label was categorized as either Ki67+ or Ki67-, based on the co-occurrence of a Ki67 label sharing the same ID. To determine the spatial distribution of cells inside the spheroid, a convex hull was constructed around the centroids of all identified cellular components within the image, effectively outlining the outer boundary of the cellular distribution. For each cell, a line was constructed, passing through the centroid of the cell and the center of the convex hull. The intersection of this line with the spheroid hull was then utilized to calculate the distance between the spheroid hull and the cell’s centroid. Scatter matrix plots were generated to visually assess the multidimensional relationships between cellular properties and marker expressions.

### 2.9. Cryosection image analysis

Images from cryosections were converted to .tif files with Fiji. The 3 most in focus z-planes were summed and used for further analysis. The spheroid’
ss area was obtained by selecting the whole spheroid in the Involucrin channel with the wand tracing tool. After median filter with radius 2, a selection was obtained for Involucrin and CK14 signals, based on thresholding. In these selections, the mean intensity and the area were measured and used to calculate the Integrated Density (mean intensity x normalized area; area was normalized to the spheroid’s area). The ratio between the Integrated Densities of the Involucrin and of the CK14 signals was calculated for each cryosection.

### 2.10. Statistical analysis

The statistical analyses in this study were conducted using GraphPad Prism 9, ensuring all data underwent tests for normal distribution. To compare the results between brightfield and 3D immunostainings, we employed an ordinary one-way ANOVA, incorporating Šidák’s correction for multiple comparisons. We established a significance threshold (α) at 0.05, corresponding to a 95% confidence interval. For the analysis of 2D immunostainings, the difference in means was assessed using Student’s t-test. In instances where pairs of groups deviated from normal distribution, we used the non-parametric Mann-Whitney U test to determine statistical significance.

## 3. Results

### 3.1. Fibroblast spheroids form in all plate types but show systemic differences in size

CCD-1137Sk human foreskin fibroblast cells were taken from freshly trypsinized adherent cultures and simultaneously seeded into 96-well plates of types A-F with 2,000 cells per well. Brightfield images were taken daily and for four days after seeding using an automated microscope. Compact, but often irregularly shaped spheroids formed in all tested plate types, and comparable amounts of loose cells were detected in all plates (Fig. 1a). On average, quantitative image analysis showed a slight decrease in spheroid diameters from day 1 to day 2 after seeding, but then, diameters remained rather stable (Fig. 1b). As anticipated from the irregular shape of most spheroids, eccentricity was ranging at a relatively high value of around 0.6 throughout the entire observation time (Fig. 1c). Eccentricity and size variations among spheroids were similar for all plate types. Conversely, there were consistent differences in spheroid diameters: First, spheroids of plate type B were smaller than those of all other types during the full experimental time window of four days (Fig. 1b, asterisks). Second, on day 4, spheroids of plate type F were larger than those of all other types (Fig. 1b, hashtag).

**Figure 1.**
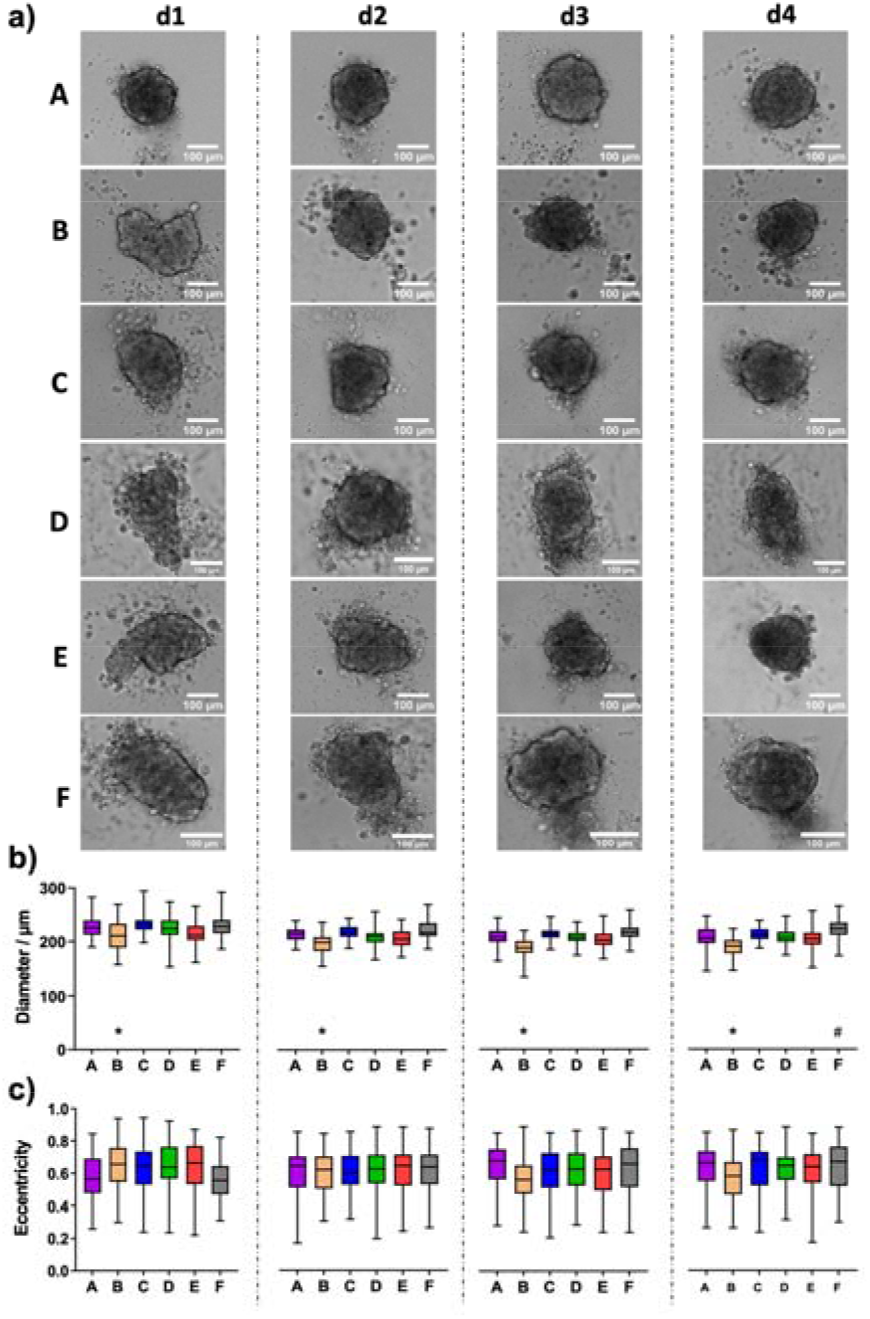
CCD-1137Sk fibroblast spheroids vary in size between different plate types. Freshly trypsinized CCD-1137Sk cells were seeded into 96-well ULA plates, types A-F, at a density of 2,000 cells per well and then cultured for up to four days. Spheroid morphology was visualized daily using automated brightfield microscopy. **a)** Representative micrographs showing individual spheroids from day 1 to 4 (d1-d4) in plate types A-F. **b-c)** Box plots depicting spheroid diameters (b) or eccentricity (c) as a function of plate type A-F. Data are from 3 experiments with ≥ 24 spheroids per experiment (mean ± SD, minima and maxima are plotted). Complete significance analysis, see Fig. S1. * and #, values significantly different compared to all other plate types for samples from the same day.

### 3.2. HaCaT spheroid size and occurrence of loose cells and satellite spheroids depends on plate type

In contrast to fibroblasts, HaCaT spheroids, grown from 5,000 freshly trypsinized cells per well, showed a continued decrease in diameter (Fig. 2a) from a value of roughly 350-380 µm on day 1 to approximately 270 µm on day 4 (Fig. 2b). Within each day of observation, the plates with the largest spheroids on day 1 continued to harbor the biggest spheroids until day 4 (Fig. 2b). From day 1 to 4, spheroids in plate type F were significantly larger than spheroids in all other types (Fig. 2b, hashtags). On days 3 and 4, spheroids in plate type A were smaller than all the others (Fig. 2b, asterisks). Since the observed variations in spheroid size may be due to different reasons, such as cell assembly, size, packaging density, or proliferation, we performed additional analyses.

**Figure 2.**
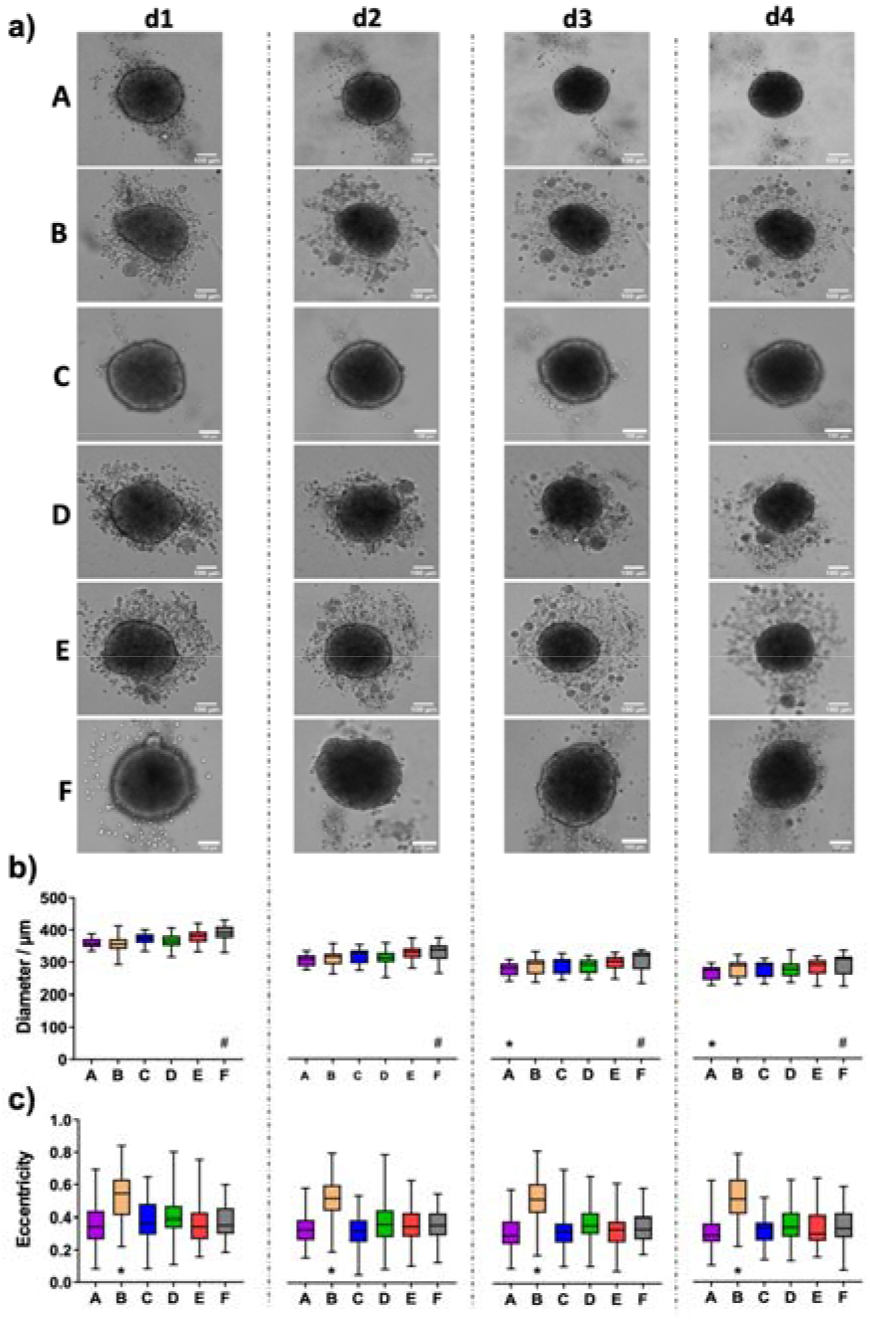
HaCaT keratinocyte spheroids vary in size and the occurrence of loose cells between different plate types. Freshly trypsinized HaCaT cells were seeded into 96-well ULA plates, types A-F, at a density of 5,000 cells per well and then cultured for up to four days. Spheroid morphology was visualized daily using automated brightfield microscopy. **a)** Representative micrographs showing individual spheroids from day 1 to day 4 (d1-d4) in plate types A-F. Scalebars, 100 µm. **b-c)** Box plots depicting spheroid diameters (b) or eccentricity (c) as a function of plate type A-F. Data are from 3 experiments with ≥ 24 spheroids per experiment (mean ± SD, minima and maxima are plotted). Complete significance analysis, see Fig. S2. ^*^ and #, values significantly different compared to all other plate types for samples from the same day.

To start with, spheroid formation is ideally characterized by the aggregation of all cells into a single and regularly shaped aggregate not surrounded by cellular satellites. While plate types A, C, and F exhibited a clear surface surrounding the spheroids, numerous loose or non-attached cells and satellite mini-spheroids were visible in all other plate types (Fig. 2a). However, this did not correlate with spheroid size, as plates A and C showed small to medium-sized spheroids. Apart from the appearance of loose cells and satellite mini-spheroids, the roundness of main spheroids varied between the different plate types. In particular, the spheroids raised in plate type B showed significantly higher eccentricity values than those of the other tested plates, meaning that their shape was less close to a perfect circle (Fig. 2c, asterisks). Over the course of four days, eccentricity values did not change significantly for any given plate type (Fig. 2c). Quantitative analysis confirmed that the area covered with loose cells and/or satellite mini-spheroids in the surroundings of the main spheroids was less than 0.5 % in plate types A and C, while it was around 4 % for types B, D, and E (Fig. 3a-b). Plate type F assumed an intermediate position with 1.4 % (Fig. 3b). As shown in Fig. 3c, the occurrence of loose cells and satellite mini-spheroids on day 4 was significantly different between plate types A, C, and F on one side, and types B, D, and E on the other side.

**Figure 3.**
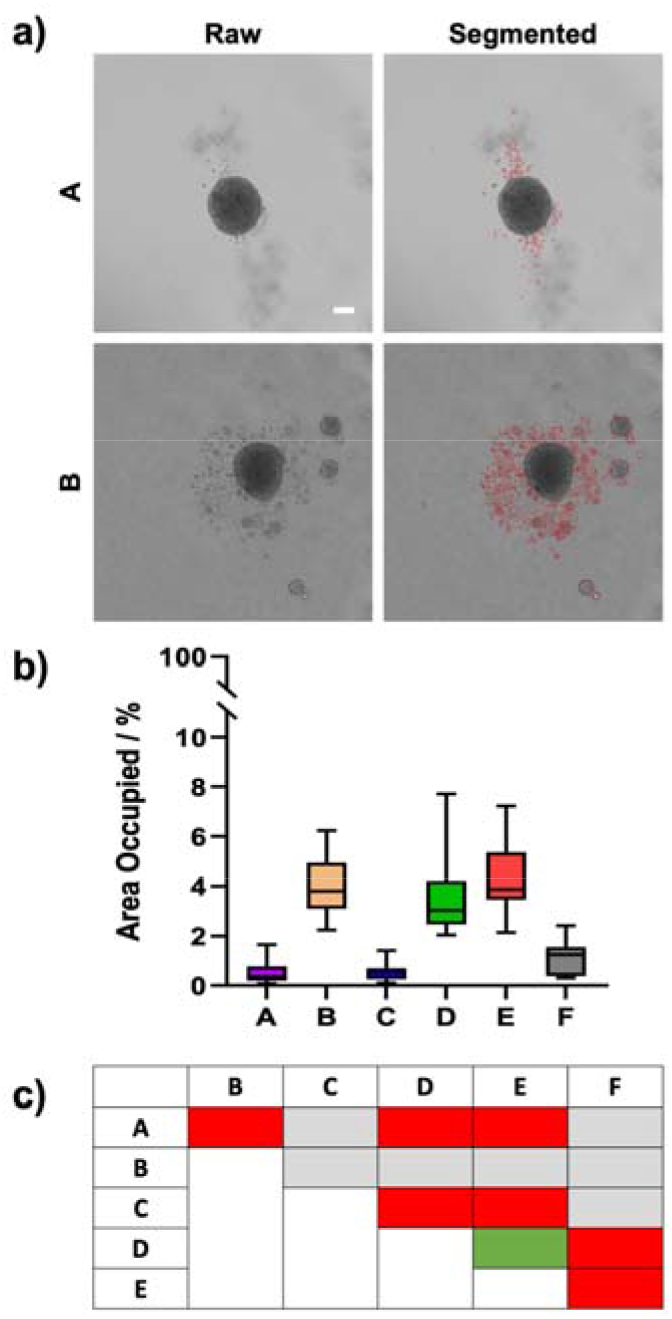
The occurrence of loose HaCaT cells and satellite mini-spheroids is dependent on plate type. Freshly trypsinized HaCaT cells were seeded into 96-well ULA plates, types A-F, at a density of 5,000 cells per well and then cultured for four days. On day 4, spheroid morphology was visualized using automated brightfield microscopy. **a**) Representative micrographs showing individual spheroids from plate types A and B, as indicated. The right panels depict regions outlined in red containing loose cells or satellite mini-spheroids that were automatically segmented from raw images (left panels). Scalebar, 100 µm. **b)** Box plot exhibiting the percentage of the image area occupied by segmented regions (as exemplified in a) as a function of plate type A-F. Data are from 3 experiments with ≥ 12 spheroids per experiment (mean ± SD, 25- and 75-percentiles are plotted). **c)** Table shows the statistical significance of Sidak multiple comparisons between the values of area occupied by loose cells and satellite mini-spheroids for plate types, as indicated. Colors stand for levels of significance: gray, n.s.; green, p ≤ 0.05; yellow, p ≤ 0.01; orange, p ≤ 0.001; red, p ≤ 0.0001.

### 3.3. Cell proliferation, differentiation and YAP1 distribution in HaCaT spheroids varies with plate type

To better understand potential reasons for the spheroid size differences, a more in-depth analysis using wholemount imaging and 3D segmentation of immunostained spheroid wholemounts from all plate types was then performed. HaCaT spheroids were fixed on day 4 and stained for nuclei, plasma membranes, the proliferation marker, Ki-67, and the proliferation- and differentiation relevant transcriptional coregulator, YAP1. After optical tissue clearing, confocal 3D-microscopy yielded full-spheroid image data stacks with a good signal-to-noise ratio throughout all samples (see Fig. 4a for representative optical sections through spheroid centers showing nuclei and Ki-67 signals). Automated AI-assisted segmentation of nuclei and Ki-67 was then performed. Consistent with the diameter analysis on live spheroids as shown in Fig. 2, spheroids from plate type A showed the smallest volumes (Fig. 4b, Fig. S3a). In addition, spheroids from plate type B were also significantly smaller than those from types C-F (Fig. 4b, Fig. S3a). Now, while the number of nuclei per spheroid was comparable between all plate types (Fig. 4c, Fig. S3b), the number of Ki-67+ cells was significantly lower in spheroids from plate type A (Fig. 4d, Fig. S3c). Further, whereas the density of nuclei packing in spheroids was only slightly different between all plate types (Fig. 4e, Fig. S3d), the distributions of nuclear volumes were significantly different between the plate types (Fig. 4f, Fig. S3e). In detail, while the nuclei of spheroids from plate types C-E were very similarly distributed, spheroids from plate types A and B showed more nuclei with smaller volumes; this was particularly prominent for nuclei from plate type A (Fig. 4f). Conversely, the size distribution of nuclei from plate type F exhibited a considerable fraction of large nuclei of around 1,000 µm^3^ (Fig. 4f).

**Figure 4.**
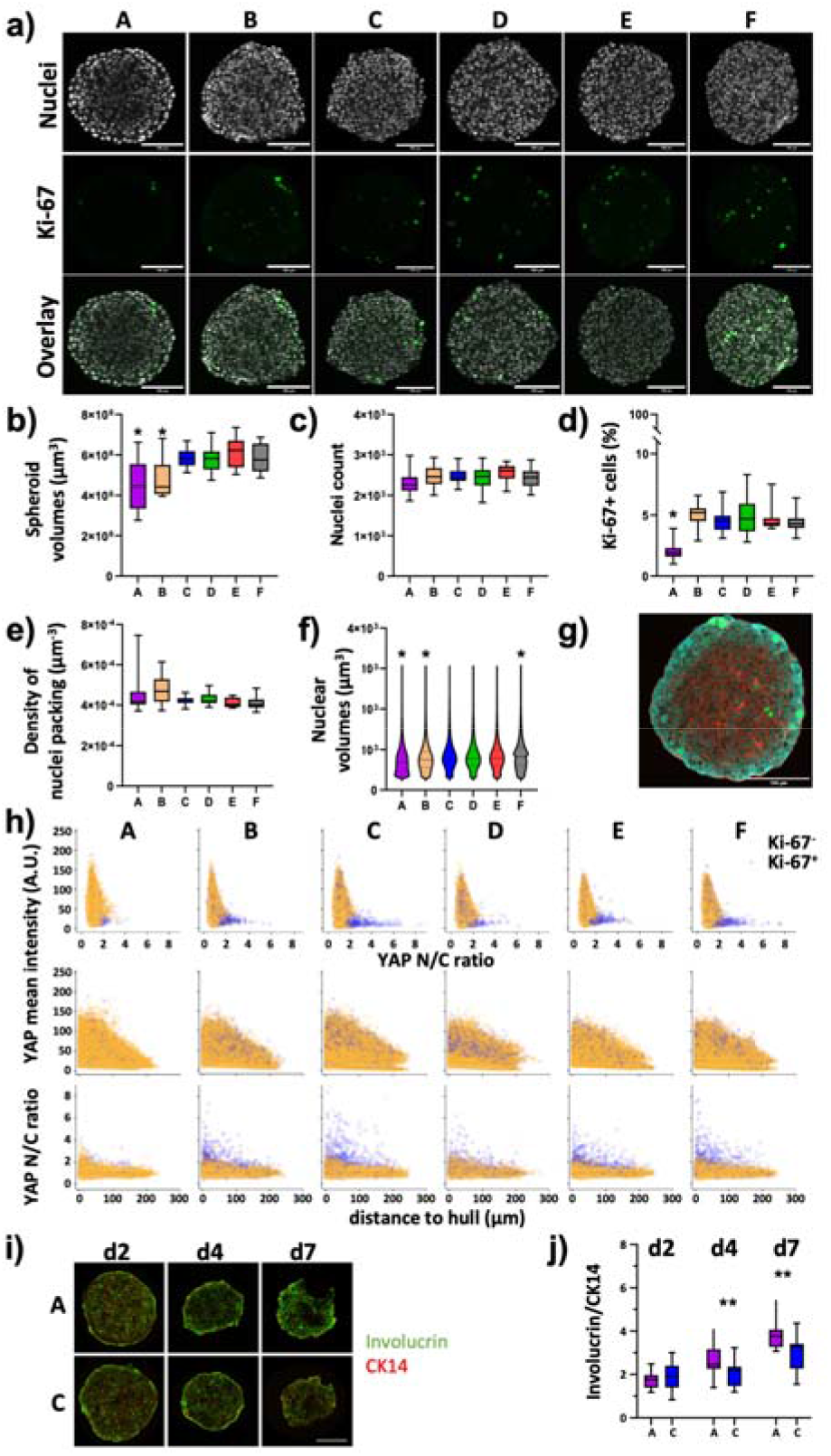
Differentiation and proliferation correlate with YAP1 in HaCaT spheroids and vary with the plate type. Freshly trypsinized HaCaT cells were seeded into 96-well ULA plates, types A-F, at a density of 5,000 cells per well and then cultured for 4 days (a-h) or for 2, 4, and 7 days (i-j). Then, spheroids were fixed, cleared, and stained for nuclei and plasma membrane, for the proliferation marker, Ki-67, and YAP1 (a-h) or fixed, cryosliced and stained for the basal keratinocyte marker, CK14, and the differenation marker, Involucrin (i-j). Wholemount confocal 3D microscopy and 3D-image segmentation (a-h) or cryosections and confocal microscopy (i-j) were performed. **a)** Representative micrographs showing single optical sections through individual spheroid wholemounts at their largest circumference, from plate types as indicated. Upper panels, DAPI nuclear signals (gray); middle panels, Ki-67 immunofluorescence signals (green); lower panels, overlays. Scalebars, 100 µm. b-e) Box plots depicting the spheroid volumes **(b)**, the number of nuclei per spheroid **(c)**, the percentage of Ki-67+ nuclei of all nuclei **(d)**, and the density of nuclei packing within spheroids **(e). (f)** Violin plot showing the size distribution of nuclear volumes in HaCaT spheroids as a function of plate type. Data in (b-f) are from 3 experiments with ≥ 24 spheroids per experiment (mean ± SD, 25- and 75-percentiles are plotted). *, values significantly different compared to all other plate types. Complete significance analysis, see Fig. S3. **(g)** Representative micrograph showing a single optical section through a spheroid wholemount at its largest circumference, from plate type F. Immunofluorescence signals of Ki-67, green; YAP1, cyan; plasma membrane, red. Scalebar, 100 µm. **(h)** Scatterplots showing values of all segmented cells (29916-42396 cells per plate type) for plate types A-F (indicated). Depicted are YAP1 mean intensity per cell as a function of YAP1 N/C ratio (upper row), YAP1 mean intensity per cell as a function of the cell’s distance to spheroid hull (middle row), YAP1 N/C ratio as a function of the cell’s distance to spheroid hull (lower row). Yellow and purple dots represent values of Ki-67- and Ki-67+ cells, respectively. **(i)** Representative confocal sum projections from cryoslices of spheroids harvested at day 2, 4, or 7 after seeding (indicated) from plate types A and C. Fluorescence signals of Involucrin, green; CK14, red. Scalebar, 100 µm. **(j)** Box plot depicting the Integrated Density ratio for Involucrin/CK14 fluorescence as a function of plate type and day of harvesting. Each data point is from 15 spheroids from three independent experiments. ^**^, p ≤ 0.01.

Previous work showed that YAP1 nuclear localization in adherent HaCaT cultures is dependent on cell density **[24]**, that YAP1 activity is primarily confined to the basal, proliferating keratinocyte layer of the epidermis **[43]**, and that YAP1 activity is increased in activated keratinocytes during wound healing **[44]**. This prompted us to investigate a potential correlation between the reduced amount of Ki-67+ HaCaT cells in spheroids from plate type A. Already at a first glance, it was evident that YAP1 signal intensity increased towards the border of the spheroids (Fig. 4g). Using the plasma membrane staining to identify the cell bodies and further algorithm pipelines to register the cell bodies to their corresponding nuclei, it became possible to automatically segment thousands of cells in these dense spheroids. This allowed to determine single-cell values of YAP1 signal intensity as well as YAP1 nucleus/cytoplasm ratio, and to co-register these values to their Ki-67 status. As depicted in Fig. 4h, this kind of analysis confirmed the qualitative impression of an increased YAP1 signal intensity towards the spheroid border in all plate types (Fig. 4h, middle row). Further, although Ki-67+ cells in plate types B-F exhibited a broad distribution of YAP1 mean intensity that largely overlapped with the values for Ki-67-cells, a subpopulation of Ki-67+ cells in these plate types exhibited a high YAP1 nucleus/cytoplasm ratio (Fig. 4h, upper row, B-F) and this ratio was increasing towards the spheroid rim (Fig. 4h, lower row, B-F). Notably, this subpopulation of Ki-67+ cells with high YAP1 nucleus/cytoplasm ratio was nearly absent in spheroids from plate type A (Fig. 4h, upper row, A), suggesting that these might be the ones missing in the total Ki-67+ cell counts (Fig. 4d).

Since keratinocyte differentiation is normally preceded by exit from cell cycle **[43]**, we asked if the reduction in proliferating cells observed in spheroids of plate type A might be due to enhanced HaCaT cell differentiation. Therefore, HaCaT spheroids were cultured in plate types A and C, harvested after 2, 4, and 7 days after seeding, cryosectioned, and stained for the basal keratinocyte marker, cytokeratin 14 (CK14), and for Involucrin, a marker of more differentiated cells. Qualitatively, this showed the expected decrease in diameter from early to late time points for spheroids from both plate types (Fig. 4i). While CK14 was evenly distributed throughout the spheroid diameters, Involucrin was more concentrated towards the spheroid borders (Fig. 4i). Quantitative analysis of the Involucrin/CK14 ratio showed that although spheroids from both plate types exhibited an increase of Involucrin/CK14 ratio with time, arguing for an ongoing differentiation process, the rise in Involucrin/CK14 ratio was more pronounced in the spheroids from plate type A (Fig. 4j). In summary, these data were consistent with a plate-type-dependent variation of YAP1 expression and its nucleus/cytoplasm ratio and with a concomittant antagonistic regulation of cell proliferation and differentiation.

### 3.3. HT-29 spheroids exhibit increasing roundness during the growth phase and slight plate-type dependent differences in size

Opposite to fibroblasts and keratinocytes, spheroids grown from 500 freshly trypsinized HT-29 colon cancer cells showed the typical, robust growth of highly proliferating cells (Fig. 4a). While HT-29 cell aggregates were still mostly irregular during days 1 and 2 after seeding, they appeared as compact spheroids on day 3 and exhibited a round shape with a sharp border on day 4, regardless of the plate type. Quantitative analysis revealed an increase in the spheroid size of roughly 26 % from day 1 to day 4 for all plate types (Fig. 5a). However, HT-29 spheroids from plate type F were consistently larger than those of all other types (Fig. 5a, Fig. S4a). Conversely, eccentricity values were largely alike between all plate types for a given day (Fig. 5b, Fig. S4b). Yet, for all plate types, eccentricity values decreased from approximately 0.5 to 0.3 (Fig. 5b), corroborating the observed increase in roundness.

**Figure 5.**
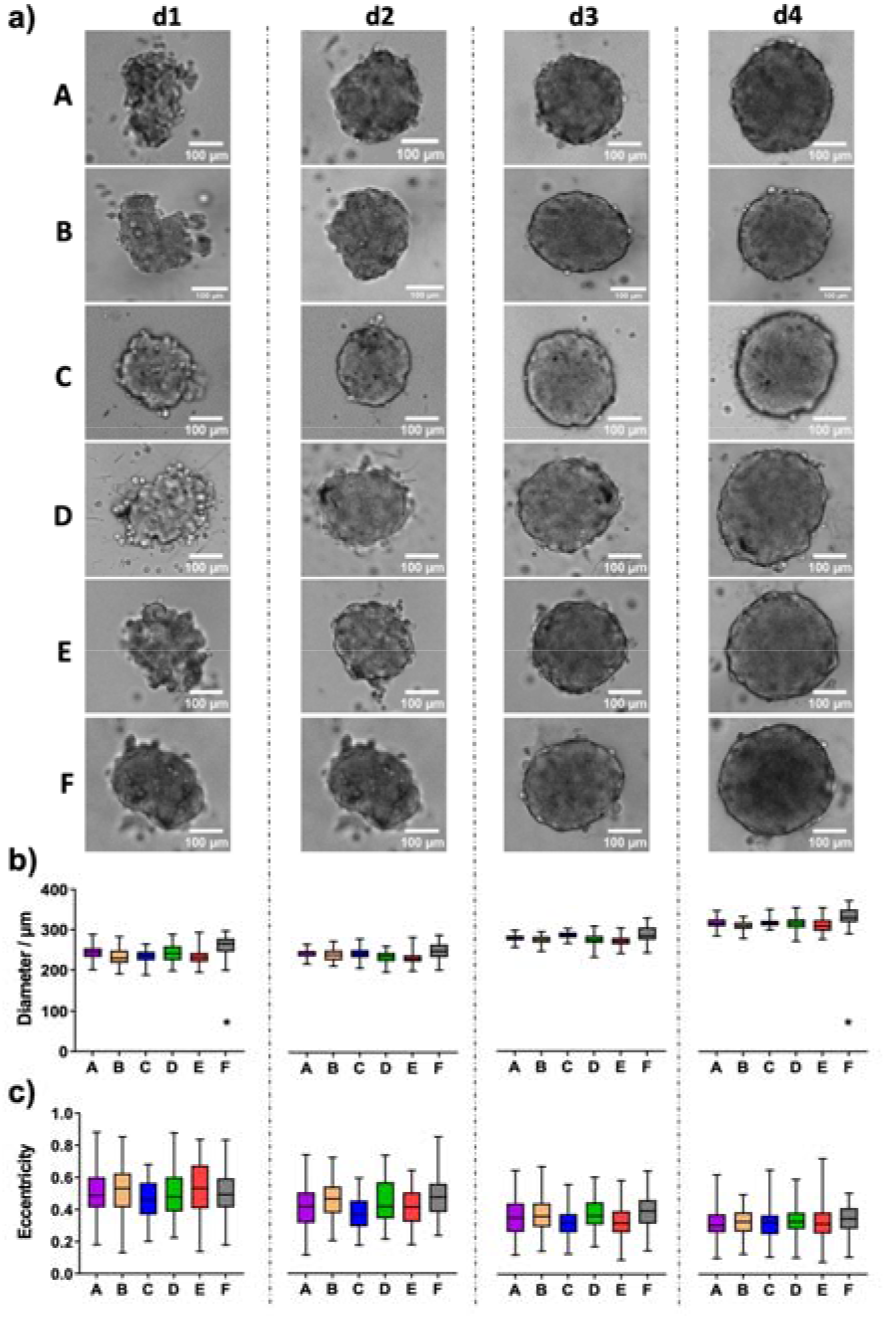
HT-29 colon cancer spheroids form robustly in all tested plate types. Freshly trypsinized HT-29 cells were seeded into 96-well ULA plates, types A-F, at a density of 500 cells per well and then cultured for up to four days. Spheroid morphology was visualized daily using automated brightfield microscopy. **a)** Representative micrographs showing individual spheroids from day 1 to day 4 (d1-d4) in plate types A-F. Scalebars, 100 µm. **b-c)** Box plots depicting spheroid diameters (b) or eccentricity (c) as a function of plate type A-F. Data are from 3 experiments with ≥ 24 spheroids per experiment (mean ± SD, 25- and 75-percentiles are plotted). Complete significance analysis, see Fig. S4. ^*^, values significantly different compared to all other plate types for samples from the same day.

Similar to the HaCaT cells, to investigate the diameter differences observed between spheroids from plate type F vs. types A-E, we again performed the wholemount confocal analysis (see Fig. 6a for representative images). Quantitative analysis of spheroid volumes (Fig. 6b) confirmed the diameter measurements in live cells (Fig. 5b). Indeed, spheroids from plate type F had larger volumes compared to all other types (Fig. 6b, Fig. S5a). This was also reflected by higher nuclei counts in plate F spheroids (Fig. 6c, Fig. S5b). Conversely, the number of proliferating cells (Fig. 6d, Fig. S5c) and the density of nuclei packing (Fig. 6e, Fig. S5d) were rather similar between all plate types. Finally, nuclei from plate type F spheroids showed a different size distribution compared to those of all other plate types (Fig. 6f, Fig. S5e). In particular, nuclei with a larger volume (> 1,000 µm^3^) were more abundant in these spheroids. Concerning YAP1, both mean intensity and N/C ratio showed a gradient from the center to the border of spheroids (Fig. 6g-h). YAP1 mean intensity of Ki-67+ cells largely overlapped with that of Ki-67-cells (Fig. 6h).

**Figure 6.**
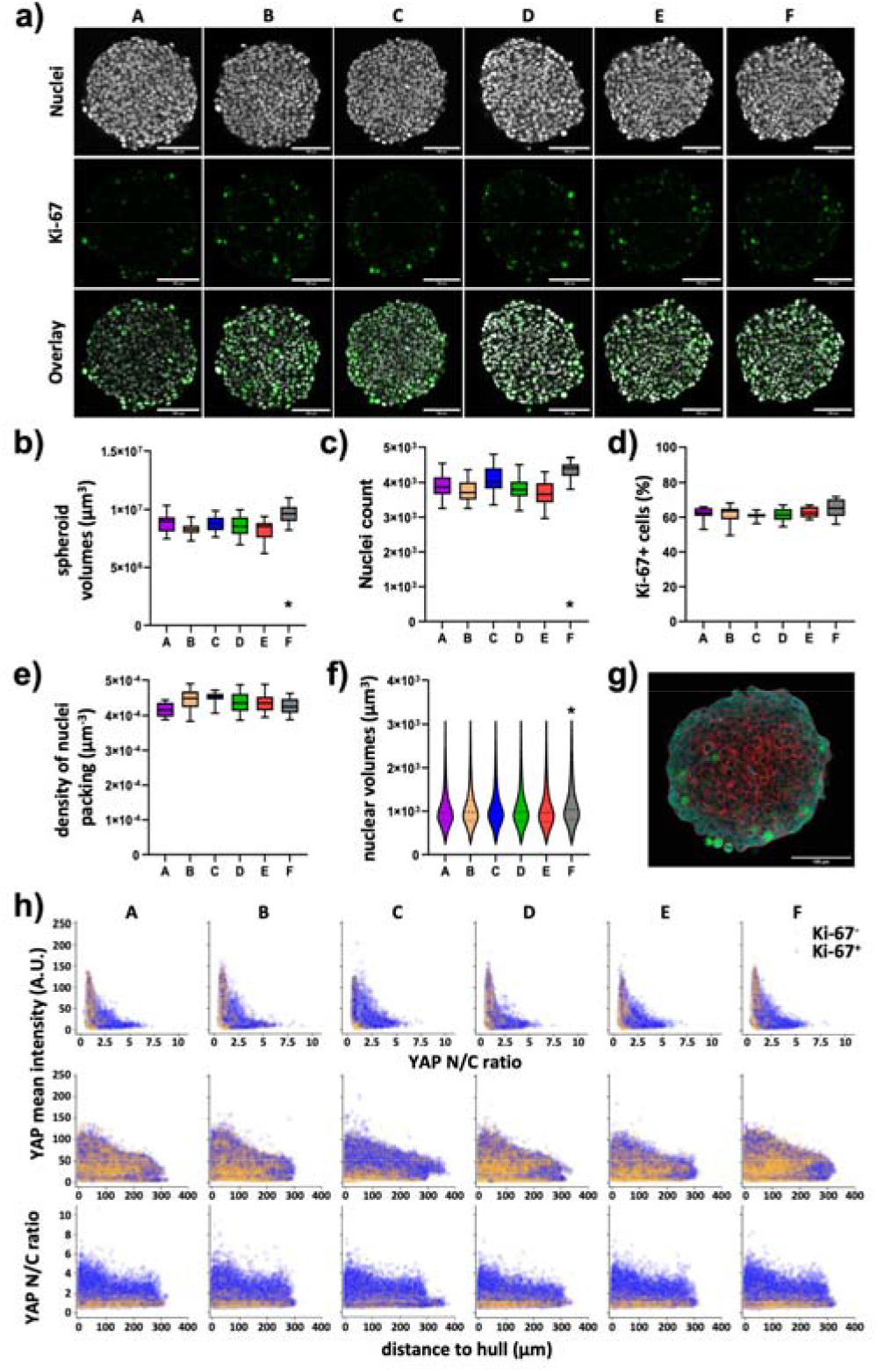
Nuclear counts and nuclear volume distribution vary in HT-29 spheroids depending on the plate type. Freshly trypsinized HT-29 cells were seeded into 96-well ULA plates, types A-F, at a density of 500 cells per well and then cultured for four days. On day 4, spheroids were fixed, cleared, and stained for nuclei and plasma membrane, for the proliferation marker, Ki-67, and YAP1. Wholemount confocal 3D microscopy and 3D-image segmentation were performed. **a)** Representative micrographs showing single optical sections through individual spheroids at their largest circumference, from plate types as indicated. Upper panels, DAPI nuclear signals (gray); middle panels, Ki-67 immunofluorescence signals (green); lower panels, overlays. Scalebars, 100 µm. b-e) Box plots depicting the spheroid volumes **(b)**, the number of nuclei per spheroid **(c)**, the percentage of Ki-67+ nuclei of all nuclei **(d)**, and the density of nuclei packing within spheroids **(e). (f)** Violin plot showing the size distribution of nuclear volumes in HT-29 spheroids as a function of plate type. Data in (b-f) are from 3 experiments with ≥ 24 spheroids per experiment (mean ± SD, 25- and 75-percentiles are plotted). *, values significantly different compared to all other plate types. Complete significance analysis, see Fig. S5. **(g)** Representative micrograph showing a single optical section through a spheroid wholemount at its largest circumference, from plate type F. Immunofluorescence signals of Ki-67, green; YAP1, cyan; plasma membrane, red. Scalebar, 100 µm. **(h)** Scatterplots showing values of all segmented cells (59,337-74,649 cells per plate type) for plate types A-F (indicated). Depicted are YAP1 mean intensity per cell as a function of YAP1 N/C ratio (upper row), YAP1 mean intensity per cell as a function of the cell’s distance to spheroid hull (middle row), YAP1 N/C ratio as a function of the cell’s distance to spheroid hull (lower row). Yellow and purple dots represent values of Ki-67- and Ki-67+ cells, respectively.

### 3.4. Formation of MDA-MB-231 spheroids is variable between plate types

Finally, we addressed the formation of spheroids using MDA-MB-231 breast cancer cells. In the presence of collagen I, the seeding of 5,000 cells per well led to a consistent generation of stable and growing spheroids for all plate types (Fig. 7a). Quantitative analysis revealed that growth from day 1 to 4 varied from 24 % to 34 % for different plate types (Fig. 7b, Fig. S6a). Furthermore, in plate types B and E, the formation of smaller satellite cell aggregates that sometimes fused to the main spheroid, was observed (Fig. 7a). This led to significantly higher eccentricity values for these plate types, however with an inconsistent pattern of the culture days (Fig. 7c, Fig. S6b).

**Figure 7.**
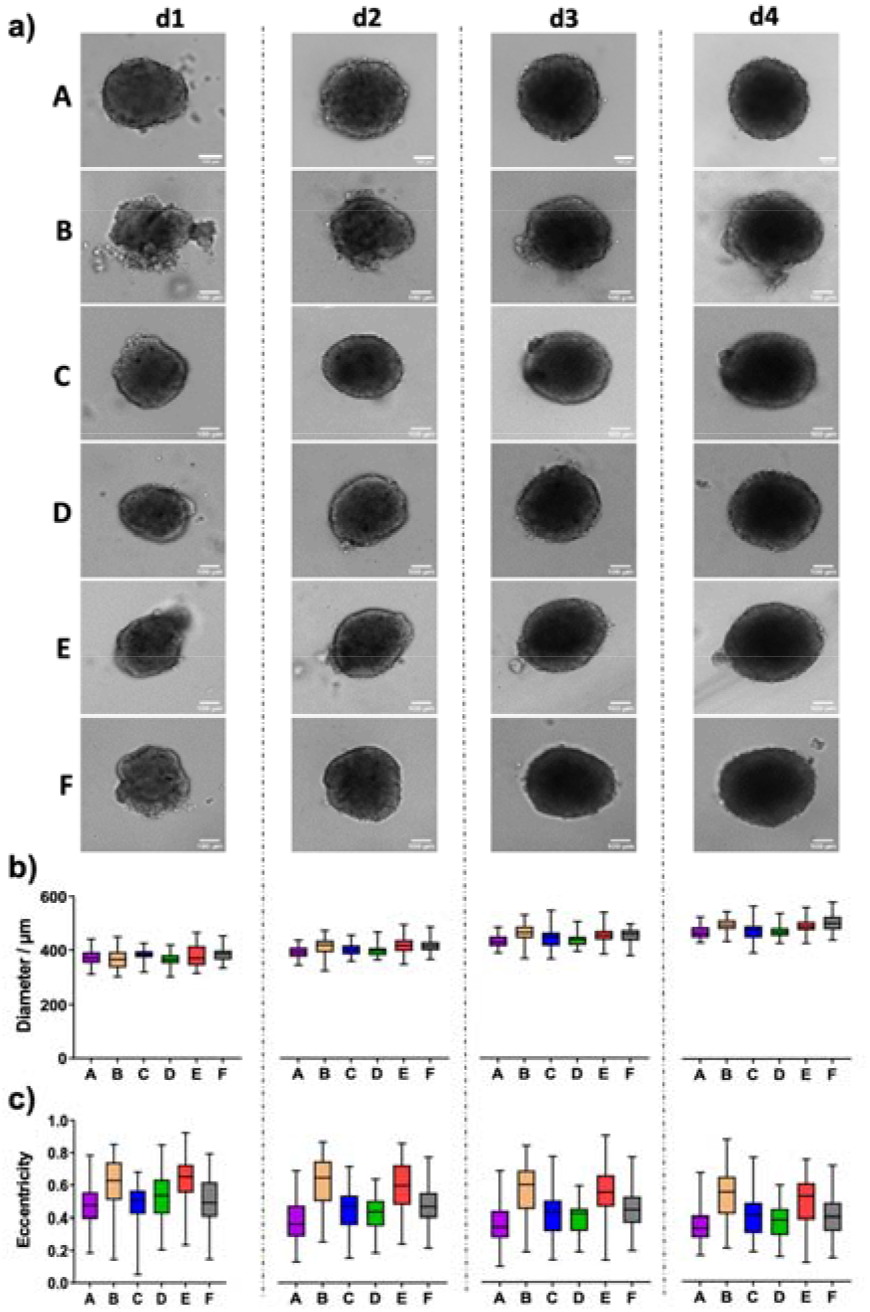
MDA-MB-231 breast cancer spheroids show satellite spheroids in plate types B and E. Freshly trypsinized MDA-MB-231 cells were seeded in the presence of 5 µg/ml collagen I into 96-well ULA plates, types A-F, at a density of 5,000 cells per well and then cultured for up to four days. Spheroid morphology was visualized daily using automated brightfield microscopy. **a)** Representative micrographs showing individual spheroids from day 1 to day 4 (d1-d4) in plate types A-F. Scalebars, 100 µm. **b-c)** Box plots depicting spheroid diameters (b) or eccentricity (c) as a function of plate type A-F. Data are from 3 experiments with ≥ 24 spheroids per experiment (mean ± SD, 25- and 75-percentiles are plotted). Complete significance analysis, see Fig. S6.

## 4. Discussion

Approaches based on three-dimensional cell cultures play a big role in fundamental and applied research, in particular, in the context of developmental and cancer biology as well as drug screening. Among the variants of three-dimensional cell cultures, spheroids serve as robust and reliable model systems with intermediate complexity and physiological relevance. Generation of spheroids can be obtained with several techniques, of which ULA plates have become increasingly important due to their ease of use and commercial availability. However, it has remained elusive, if and to what extent different ULA-plate types affect the outcome of spheroid formation and growth. For six different plate types, the present study systematically addressed morphological characteristics of spheroids prepared from four distinct cell types, over the course of four days after seeding. Although this revealed that, in principle, all tested plate types succeeded in reliably producing spheroids of all cell lines under investigation, significant differences in spheroid size and roundness, as well as cell number and proliferation were observed in a cell-line-dependent manner. This highlights that the choice of plate type may affect experimental outcomes and that consistency in the use of a specific plate type is critical during experimental campaigns.

As a major morphological parameter, we systematically measured the size of spheroids during an experimental time course of four days after seeding. As expected, growth was negative or static for non-neoplastic fibroblast cells and keratinocytes, while it was positive for colon and breast cancer cells. These patterns were similar between all six ULA-plate types tested. However, spheroids of all cell types raised in plate type F were consistently larger than in all other plate types and those from plate types A (for HaCaT and MDA-MB-231) and B (for CCD-1137Sk and HaCaT) were smaller than spheroids from the other plate types.

A more in-depth confocal wholemount analysis for HaCaT and HT-29 cells revealed that the reasons for these discrepancies might be different ones, depending on the cell type. Indeed, for HaCaT cells, the overall cell number was similar when comparing spheroids from all six plate types, while it was significantly different in the case of HT-29 cells. Thus, while for HT-29 cells a higher cell number might explain the larger spheroids in plate type F, this scenario was unlikely in HaCaT spheroids. To investigate this further, nuclear volume was determined as a putative reason for the observed differences in the spheroid volume of HaCaT spheroids. Indeed, while the HaCaT spheroids from plates A and B were small and had smaller nuclei than spheroids from other plates, the largest HaCaT spheroids, which were from plate type F, also showed the largest nuclei, and HaCaT spheroids with intermediate volumes from plates C-E also showed intermediate nuclei volumes.

In general, nuclear size and shape are determined by several factors, including alterations in nuclear transport, cell cycle regulation, genome packaging, genome activity, cellular physiology, and/or cell viability **[45]**. In the specific case of HaCaT keratinocytes, nuclear morphology varied according to their differentiation status **[46]**, at least in 2D cultures, where incubation in high Ca^2+^ levels leads to an increased expression of differentiated keratinocyte markers **[47]**. In spheroids, HaCaT cells were also found to exhibit a stratification with more differentiated cells towards the spheroid rim **[20]**. Furthermore, during epidermal differentiation, keratinocytes become postmitotic before they go into apoptosis [48]. Thus, as for many epithelial cell types, there is an inverse correlation between proliferation and differentiation of epidermal keratinocytes. The Hippo signaling pathway with its downstream effectors YAP and TAZ is particularly relevant to control epidermal homeostasis. Indeed, active, i.e., nuclear YAP is normally present only in keratinocytes showing proliferative activity, namely in the basal layer of the epidermis **[43]**. Furthermore, it is upregulated in states of enhanced growth, i.e., upon wound healing and in epidermal cancers **[43]**. This was corroborated by our findings that HaCaT spheroids from plate type A showed a lower amount of Ki-67+ proliferating cells (Fig. S7), a higher propensity to maturate and a lack Ki-67+ exhibiting a high YAP1 N/C ratio. Regarding the HT-29 colon cancer cells, only spheroids grown in plate type F were significantly different in size from those raised in all other plate types. As mentioned earlier, this could be likely explained by the higher cell numbers counted in HT-29 spheroids from plate type F. However, in these samples, the nuclear volumes were significantly larger than in those from plate types A-E. Currently, it is unclear if this contributed to the observed differences in spheroid size. Moreover, the underlying reasons for the altered nuclear volume are difficult to explain. Indeed, cell proliferation as evaluated by the fraction of Ki-67+ cells as well as the analysis of YAP1 distribution and N/C ratio were inconspicuous (Fig. S7). Further options would be differences in apoptosis, pressure [49], or stiffness.

Finally, this study also revealed plate-type dependent differences in spheroid roundness and the propensity of leaving loose cells or satellite spheroids. These effects were primarily observed in HaCaT and MDA-MB-231 cells. Since details on the surfaces of ULA plates are usually not disclosed, it is unclear what induced the detected differences. However, it has become clear that the selection of a specific plate type can have profound consequences on major morphological and cell biological parameters. This warrants careful consideration upfront to the execution of experimental campaigns and supports the need of implementing different plate types depending on the nature of the investigation.

## Author Contributions

Conceptualization, M.V. and R.R.; methodology, M.V., A.A., R.B., T.C., F.K.; software, M.V., R.B., E.N., F.P., M.R., K.S.; validation, M.V., A.A.; formal analysis, M.V., A.A., T.C., E.N.; investigation, A.A., T.C., F.K.; resources, M.R., M.H., R.R.; data curation, M.V., A.A.; writing—original draft preparation, M.V., A.A., R.R.; writing—review and editing, M.V., R.R.; visualization, M.V., A.A.; supervision, M.R., M.H., R.R.; project administration, M.V., R.R.; funding acquisition, M.R., S.S., M.H., R.R.. All authors have read and agreed to the published version of the manuscript.

## Funding

This work was funded by the German Federal Ministry of Education and Research (BMBF) grant 01IS21062B. RB and RR were funded by the Carl-Zeiss Foundation, project DigiFIT. This work was funded by the German Federal Ministry of Education and Research (BMBF) as part of the Innovation Partnership M^2^Aind, projects Drugs4Future (13FH8I05IA) and DrugsData (13FH8I09IA) within the framework Starke Fachhochschulen-Impuls für die Region (FH-Impuls). This research project is part of the Forschungscampus M^2^OLIE and funded by the German Federal Ministry of Education and Research (BMBF) within the “Framework Forschungscampus: public-private partnership for Innovations”. This work was supported by DFG grant INST874/9-1. This work was partially funded by faCellitate GmbH. This work was supported by the Human Frontier Science Program (career development award to K.M.S.) and the Helmholtz Gesellschaft.

## Data Availability Statement

Data available on request.

## Conflicts of Interest

The funders had no role in the design of the study; in the collection, analyses, or interpretation of data; in the writing of the manuscript; or in the decision to publish the results.

## Notes

### Competing Interest Statement

The authors have declared no competing interest.

### Summary of Updates

Supplements were added. Further, we have reformatted the manuscript. We have deleted the format of the target journal.

