## Supplement for "A Multiparametric Analysis Reveals Differential Behavior of Spheroid Cultures on Distinct Ultra-Low Attachment Plates Types"

**a)**

|  | B | | | | C | | | | D | | | | E | | | | F | | | |
| --- | --- | --- | --- | --- | --- | --- | --- | --- | --- | --- | --- | --- | --- | --- | --- | --- | --- | --- | --- | --- |
|  | d1 | d2 | d3 | d4 | d1 | d2 | d3 | d4 | d1 | d2 | d3 | d4 | d1 | d2 | d3 | d4 | d1 | d2 | d3 | d4 |
| A |  |  |  |  |  |  |  |  |  |  |  |  |  |  |  |  |  |  |  |  |
| B |  | | | |  |  |  |  |  |  |  |  |  |  |  |  |  |  |  |  |
| C |  |  |  |  |  | | | |  |  |  |  |  |  |  |  |  |  |  |  |
| D |  |  |  |  |  |  |  |  |  | | | |  |  |  |  |  |  |  |  |
| E |  |  |  |  |  |  |  |  |  |  |  |  |  | | | |  |  |  |  |

**b)**

|  | B | | | | C | | | | D | | | | E | | | | F | | | |
| --- | --- | --- | --- | --- | --- | --- | --- | --- | --- | --- | --- | --- | --- | --- | --- | --- | --- | --- | --- | --- |
|  | d1 | d2 | d3 | d4 | d1 | d2 | d3 | d4 | d1 | d2 | d3 | d4 | d1 | d2 | d3 | d4 | d1 | d2 | d3 | d4 |
| A |  |  |  |  |  |  |  |  |  |  |  |  |  |  |  |  |  |  |  |  |
| B |  | | | |  |  |  |  |  |  |  |  |  |  |  |  |  |  |  |  |
| C |  |  |  |  |  | | | |  |  |  |  |  |  |  |  |  |  |  |  |
| D |  |  |  |  |  |  |  |  |  | | | |  |  |  |  |  |  |  |  |
| E |  |  |  |  |  |  |  |  |  |  |  |  |  | | | |  |  |  |  |

**Figure S1:** ***CCD-1137Sk spheroids vary primarily in size between different plate types.*** Tables display statistical significance of Sidak multiple comparisons between values of **a)** spheroid diameter and **b)** spheroid eccentricity for plate types and days in culture (d1-d4) as indicated, experimental data as depicted in Figure 1. Colors indicate levels of significance: gray, n.s.; green, p ≤ 0.05; yellow, p ≤ 0.01; orange, p ≤ 0.001; red, p ≤ 0.0001.

**a)**

|  | B | | | | C | | | | D | | | | E | | | | F | | | |
| --- | --- | --- | --- | --- | --- | --- | --- | --- | --- | --- | --- | --- | --- | --- | --- | --- | --- | --- | --- | --- |
|  | d1 | d2 | d3 | d4 | d1 | d2 | d3 | d4 | d1 | d2 | d3 | d4 | d1 | d2 | d3 | d4 | d1 | d2 | d3 | d4 |
| A |  |  |  |  |  |  |  |  |  |  |  |  |  |  |  |  |  |  |  |  |
| B |  | | | |  |  |  |  |  |  |  |  |  |  |  |  |  |  |  |  |
| C |  |  |  |  |  | | | |  |  |  |  |  |  |  |  |  |  |  |  |
| D |  |  |  |  |  |  |  |  |  | | | |  |  |  |  |  |  |  |  |
| E |  |  |  |  |  |  |  |  |  |  |  |  |  | | | |  |  |  |  |

**b)**

|  | B | | | | C | | | | D | | | | E | | | | F | | | |
| --- | --- | --- | --- | --- | --- | --- | --- | --- | --- | --- | --- | --- | --- | --- | --- | --- | --- | --- | --- | --- |
|  | d1 | d2 | d3 | d4 | d1 | d2 | d3 | d4 | d1 | d2 | d3 | d4 | d1 | d2 | d3 | d4 | d1 | d2 | d3 | d4 |
| A |  |  |  |  |  |  |  |  |  |  |  |  |  |  |  |  |  |  |  |  |
| B |  | | | |  |  |  |  |  |  |  |  |  |  |  |  |  |  |  |  |
| C |  |  |  |  |  | | | |  |  |  |  |  |  |  |  |  |  |  |  |
| D |  |  |  |  |  |  |  |  |  | | | |  |  |  |  |  |  |  |  |
| E |  |  |  |  |  |  |  |  |  |  |  |  |  | | | |  |  |  |  |

**Figure S2:** ***HaCaT spheroid size and eccentricity depend on plate type.*** Tables display statistical significance of Sidak multiple comparisons between values of **a)** spheroid diameter and **b)** spheroid eccentricity for plate types and days in culture (d1-d4) as indicated, experimental data as depicted in Figure 2. Colors indicate levels of significance: gray, n.s.; green, p ≤ 0.05; yellow, p ≤ 0.01; orange, p ≤ 0.001; red, p ≤ 0.0001.

**a)**

|  | B | C | D | E | F |
| --- | --- | --- | --- | --- | --- |
| A |  |  |  |  |  |
| B |  |  |  |  |  |
| C |  |  |  |  |  |
| D |  |  |  |  |  |
| E |  |  |  |  |  |

**b)**

|  | B | C | D | E | F |
| --- | --- | --- | --- | --- | --- |
| A |  |  |  |  |  |
| B |  |  |  |  |  |
| C |  |  |  |  |  |
| D |  |  |  |  |  |
| E |  |  |  |  |  |

**c)**

|  | B | C | D | E | F |
| --- | --- | --- | --- | --- | --- |
| A |  |  |  |  |  |
| B |  |  |  |  |  |
| C |  |  |  |  |  |
| D |  |  |  |  |  |
| E |  |  |  |  |  |

**d)**

|  | B | C | D | E | F |
| --- | --- | --- | --- | --- | --- |
| A |  |  |  |  |  |
| B |  |  |  |  |  |
| C |  |  |  |  |  |
| D |  |  |  |  |  |
| E |  |  |  |  |  |

**e)**

|  | B | C | D | E | F |
| --- | --- | --- | --- | --- | --- |
| A |  |  |  |  |  |
| B |  |  |  |  |  |
| C |  |  |  |  |  |
| D |  |  |  |  |  |
| E |  |  |  |  |  |

**Figure S3:** ***HaCaT spheroids primarily vary in frequency of Ki67+ cells, in volume, and in nuclear volume between different plate types.*** Tables display statistical significance of Sidak multiple comparisons between values of **a)** spheroid volume, **b)** nuclear count per spheroid, **c)** fraction of Ki67+ nuclei, **d)** density of nuclei packing, and **e)** volume of individual nuclei for plate types as indicated, experimental data as depicted in Figure 4. Colors indicate levels of significance: gray, n.s.; green, p ≤ 0.05; yellow, p ≤ 0.01; orange, p ≤ 0.001; red, p ≤ 0.0001.

**a)**

|  | B | | | | C | | | | D | | | | E | | | | F | | | |
| --- | --- | --- | --- | --- | --- | --- | --- | --- | --- | --- | --- | --- | --- | --- | --- | --- | --- | --- | --- | --- |
|  | d1 | d2 | d3 | d4 | d1 | d2 | d3 | d4 | d1 | d2 | d3 | d4 | d1 | d2 | d3 | d4 | d1 | d2 | d3 | d4 |
| A |  |  |  |  |  |  |  |  |  |  |  |  |  |  |  |  |  |  |  |  |
| B |  | | | |  |  |  |  |  |  |  |  |  |  |  |  |  |  |  |  |
| C |  |  |  |  |  | | | |  |  |  |  |  |  |  |  |  |  |  |  |
| D |  |  |  |  |  |  |  |  |  | | | |  |  |  |  |  |  |  |  |
| E |  |  |  |  |  |  |  |  |  |  |  |  |  | | | |  |  |  |  |

**b)**

|  | B | | | | C | | | | D | | | | E | | | | F | | | |
| --- | --- | --- | --- | --- | --- | --- | --- | --- | --- | --- | --- | --- | --- | --- | --- | --- | --- | --- | --- | --- |
|  | d1 | d2 | d3 | d4 | d1 | d2 | d3 | d4 | d1 | d2 | d3 | d4 | d1 | d2 | d3 | d4 | d1 | d2 | d3 | d4 |
| A |  |  |  |  |  |  |  |  |  |  |  |  |  |  |  |  |  |  |  |  |
| B |  | | | |  |  |  |  |  |  |  |  |  |  |  |  |  |  |  |  |
| C |  |  |  |  |  | | | |  |  |  |  |  |  |  |  |  |  |  |  |
| D |  |  |  |  |  |  |  |  |  | | | |  |  |  |  |  |  |  |  |
| E |  |  |  |  |  |  |  |  |  |  |  |  |  | | | |  |  |  |  |

**Figure S4:** ***HT-29 spheroid eccentricity is largely similar between different plate types, but spheroid diameter may vary.*** Tables display statistical significance of Sidak multiple comparisons between values of **a)** spheroid diameter and **b)** spheroid eccentricity for plate types and days in culture (d1-d4) as indicated, experimental data as depicted in Figure 4. Colors indicate levels of significance: gray, n.s.; green, p ≤ 0.05; yellow, p ≤ 0.01; orange, p ≤ 0.001; red, p ≤ 0.0001.

**a)**

|  | B | C | D | E | F |
| --- | --- | --- | --- | --- | --- |
| A |  |  |  |  |  |
| B |  |  |  |  |  |
| C |  |  |  |  |  |
| D |  |  |  |  |  |
| E |  |  |  |  |  |

**b)**

|  | B | C | D | E | F |
| --- | --- | --- | --- | --- | --- |
| A |  |  |  |  |  |
| B |  |  |  |  |  |
| C |  |  |  |  |  |
| D |  |  |  |  |  |
| E |  |  |  |  |  |

**c)**

|  | B | C | D | E | F |
| --- | --- | --- | --- | --- | --- |
| A |  |  |  |  |  |
| B |  |  |  |  |  |
| C |  |  |  |  |  |
| D |  |  |  |  |  |
| E |  |  |  |  |  |

**d)**

|  | B | C | D | E | F |
| --- | --- | --- | --- | --- | --- |
| A |  |  |  |  |  |
| B |  |  |  |  |  |
| C |  |  |  |  |  |
| D |  |  |  |  |  |
| E |  |  |  |  |  |

**e)**

|  | B | C | D | E | F |
| --- | --- | --- | --- | --- | --- |
| A |  |  |  |  |  |
| B |  |  |  |  |  |
| C |  |  |  |  |  |
| D |  |  |  |  |  |
| E |  |  |  |  |  |

**Figure S5:** ***HT-29 spheroids primarily vary in nuclear counts, spheroid volume, and nuclear volume between different plate types.*** Tables display statistical significance of Sidak multiple comparisons between values of **a)** spheroid volume, **b)** nuclear count per spheroid, **c)** fraction of Ki67+ nuclei, **d)** density of nuclei packing, and **e)** volume of individual nuclei for plate types as indicated, experimental data as depicted in Figure 7. Colors indicate levels of significance: gray, n.s.; green, p ≤ 0.05; yellow, p ≤ 0.01; orange, p ≤ 0.001; red, p ≤ 0.0001.

**a)**

|  | B | | | | C | | | | D | | | | E | | | | F | | | |
| --- | --- | --- | --- | --- | --- | --- | --- | --- | --- | --- | --- | --- | --- | --- | --- | --- | --- | --- | --- | --- |
|  | d1 | d2 | d3 | d4 | d1 | d2 | d3 | d4 | d1 | d2 | d3 | d4 | d1 | d2 | d3 | d4 | d1 | d2 | d3 | d4 |
| A |  |  |  |  |  |  |  |  |  |  |  |  |  |  |  |  |  |  |  |  |
| B |  | | | |  |  |  |  |  |  |  |  |  |  |  |  |  |  |  |  |
| C |  |  |  |  |  | | | |  |  |  |  |  |  |  |  |  |  |  |  |
| D |  |  |  |  |  |  |  |  |  | | | |  |  |  |  |  |  |  |  |
| E |  |  |  |  |  |  |  |  |  |  |  |  |  | | | |  |  |  |  |

**b)**

|  | B | | | | C | | | | D | | | | E | | | | F | | | |
| --- | --- | --- | --- | --- | --- | --- | --- | --- | --- | --- | --- | --- | --- | --- | --- | --- | --- | --- | --- | --- |
|  | d1 | d2 | d3 | d4 | d1 | d2 | d3 | d4 | d1 | d2 | d3 | d4 | d1 | d2 | d3 | d4 | d1 | d2 | d3 | d4 |
| A |  |  |  |  |  |  |  |  |  |  |  |  |  |  |  |  |  |  |  |  |
| B |  | | | |  |  |  |  |  |  |  |  |  |  |  |  |  |  |  |  |
| C |  |  |  |  |  | | | |  |  |  |  |  |  |  |  |  |  |  |  |
| D |  |  |  |  |  |  |  |  |  | | | |  |  |  |  |  |  |  |  |
| E |  |  |  |  |  |  |  |  |  |  |  |  |  | | | |  |  |  |  |

**Figure S6:** ***MDA-MB-231 spheroid generation shows several distinctions between different plate types.*** Tables display statistical significance of Sidak multiple comparisons between values of **a)** spheroid diameter and **b)** spheroid eccentricity for plate types and days in culture (d1-d4) as indicated, experimental data as depicted in Figure 5. Colors indicate levels of significance: gray, n.s.; green, p ≤ 0.05; yellow, p ≤ 0.01; orange, p ≤ 0.001; red, p ≤ 0.0001.

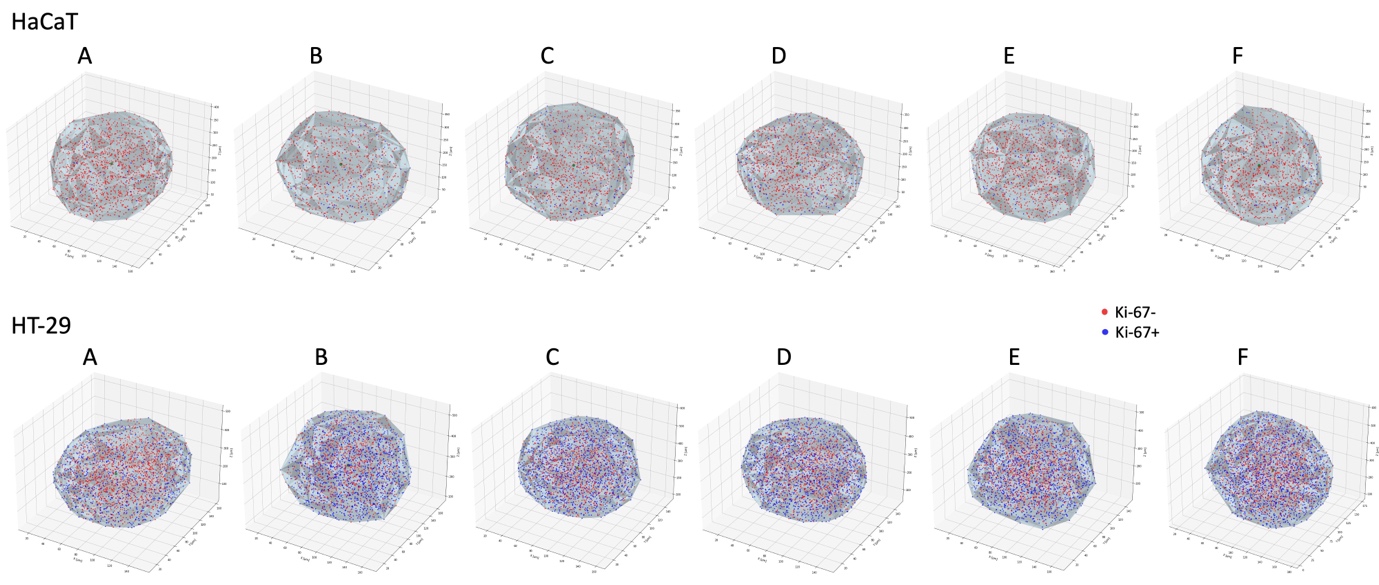

**Figure S7:** ***Frequency of Ki-67+ cells varies between HaCaT and HT-29 cells as well as between different plate types.*** 3D-plots show images of the distribution of Ki-67- (red dots) and Ki-67+ cells (blue dots) within individual representative spheroids. Grey hull shows the outline of spheroids, the center of mass of the spheroids is indicated by a green dot.
